# *In vivo* Auto-tuning of Antibody-Drug Conjugate Delivery for Effective Immunotherapy using High-Avidity, Low-Affinity Antibodies

**DOI:** 10.1101/2024.04.06.588433

**Authors:** Anna Kopp, Shujun Dong, Hyeyoung Kwon, Tiexin Wang, Alec A. Desai, Jennifer J. Linderman, Peter Tessier, Greg M. Thurber

## Abstract

Antibody-drug conjugates (ADCs) have experienced a surge in clinical approvals in the past five years. Despite this success, a major limitation to ADC efficacy in solid tumors is poor tumor penetration, which leaves many cancer cells untargeted. Increasing antibody doses or co-administering ADC with an unconjugated antibody can improve tumor penetration and increase efficacy when target receptor expression is high. However, it can also reduce efficacy in low-expression tumors where ADC delivery is limited by cellular uptake. This creates an intrinsic problem because many patients express different levels of target between tumors and even within the same tumor. Here, we generated High-Avidity, Low-Affinity (HALA) antibodies that can automatically tune the cellular ADC delivery to match the local expression level. Using HER2 ADCs as a model, HALA antibodies were identified with the desired HER2 expression-dependent competitive binding with ADCs *in vitro*. Multi-scale distribution of trastuzumab emtansine and trastuzumab deruxtecan co-administered with the HALA antibody were analyzed *in vivo*, revealing that the HALA antibody increased ADC tumor penetration in high-expression systems with minimal reduction in ADC uptake in low-expression tumors. This translated to greater ADC efficacy in immunodeficient mouse models across a range of HER2 expression levels. Furthermore, Fc-enhanced HALA antibodies showed improved Fc-effector function at both high and low expression levels and elicited a strong response in an immunocompetent mouse model. These results demonstrate that HALA antibodies can expand treatment ranges beyond high expression targets and leverage strong immune responses.

## Introduction

Antibody-drug conjugates (ADCs) combine potent cytotoxic payloads with monoclonal antibodies that bind tumor-associated cancer antigens, thereby concentrating the payload within the tumor to increase the therapeutic window. ADCs have experienced a recent surge in clinical approvals, with seven drugs approved in the past five years including several for solid tumors. One of the major limitations of this drug class in solid tumors is limited tissue penetration, leaving many cancer cells with no treatment.^1–5^ The newest generation of ADCs in solid tumors partly rely on higher antibody dosing enabled by reduced potency, such as Trodelvy (10 mg/kg twice every three weeks) and T-Dxd (5.4 or 6.4 mg/kg every three weeks), which can help overcome limited tissue penetration.^2^ Due to their large molecular weight, ADCs and other antibody-based biologics experience slow diffusion through tissues, but rapid binding to the target antigen immobilizes the ADC near blood vessels, creating the “binding site barrier.”^4,6^ The heterogeneous distribution of antibodies and ADCs has been documented extensively in preclinical models^1–3,7^ and, more recently, these trends have also been demonstrated in clinical tumors.^8,9^

There are multiple approaches to overcome this perivascular saturation front, including small molecule formats,^10,11^ reduced affinity,^12^ and paratope blocking.^13^ We have explored the use of co-administering unconjugated “carrier” antibody to partially block ADC binding, thereby driving the ADC deeper into the tissue. One advantage of this approach is the additional antibody dose that can leverage antibody-intrinsic mechanisms of action, such as receptor signaling blockade and/or Fc-effector function.^14,15^ The use of a carrier dose results in a greater number of tumor cells receiving treatment, and several high-expression models have illustrated improved efficacy with this approach.^16–18^ Importantly, the amount of drug delivered to cells on average should be tailored to the potency of the payload to improve efficacy. Thus, a ratio of carrier dose to ADC that is too high can reduce efficacy by lowering ADC delivery to cells below a lethal threshold.^19^ Similarly, the use of a carrier dose in low-expression systems, where there are fewer receptors to internalize the ADC, has shown mixed results on ADC efficacy.^15,17^ This creates a problem because patients can have variable expression between lesions (e.g. the primary tumor and metastases) and within the same lesion.^21,22^ Therefore, the optimal carrier dose does not solely rely on the average expression for a given patient.

To overcome this limitation, we aimed to improve the carrier dose approach by using an antibody that selectively competes with ADC binding on high expression cells, maximizing tissue penetration and efficacy in these regions, but does not compete on low expression cells, maximizing ADC uptake and efficacy for these cells. The concept of High-Avidity, Low-Affinity (HALA) antibodies to achieve this goal has been explored theoretically^23^ and could be used to expand the use of carrier doses to benefit a wide range of expression levels. HALA antibodies rely on differences in avidity (modeled previously^24–27^), allowing the HALA antibody to compete with the high-affinity ADC to drive better tissue penetration and efficacy on high-expression cells (**Figure 1A-1B**) while preventing blocking and maximizing ADC uptake on low-expression cells (**Figure 1C-1D**). Therefore, the HALA and ADC mixture ‘auto-tunes’ the delivery of ADC to each cell based on the most effective ratio.

**Figure 1.**
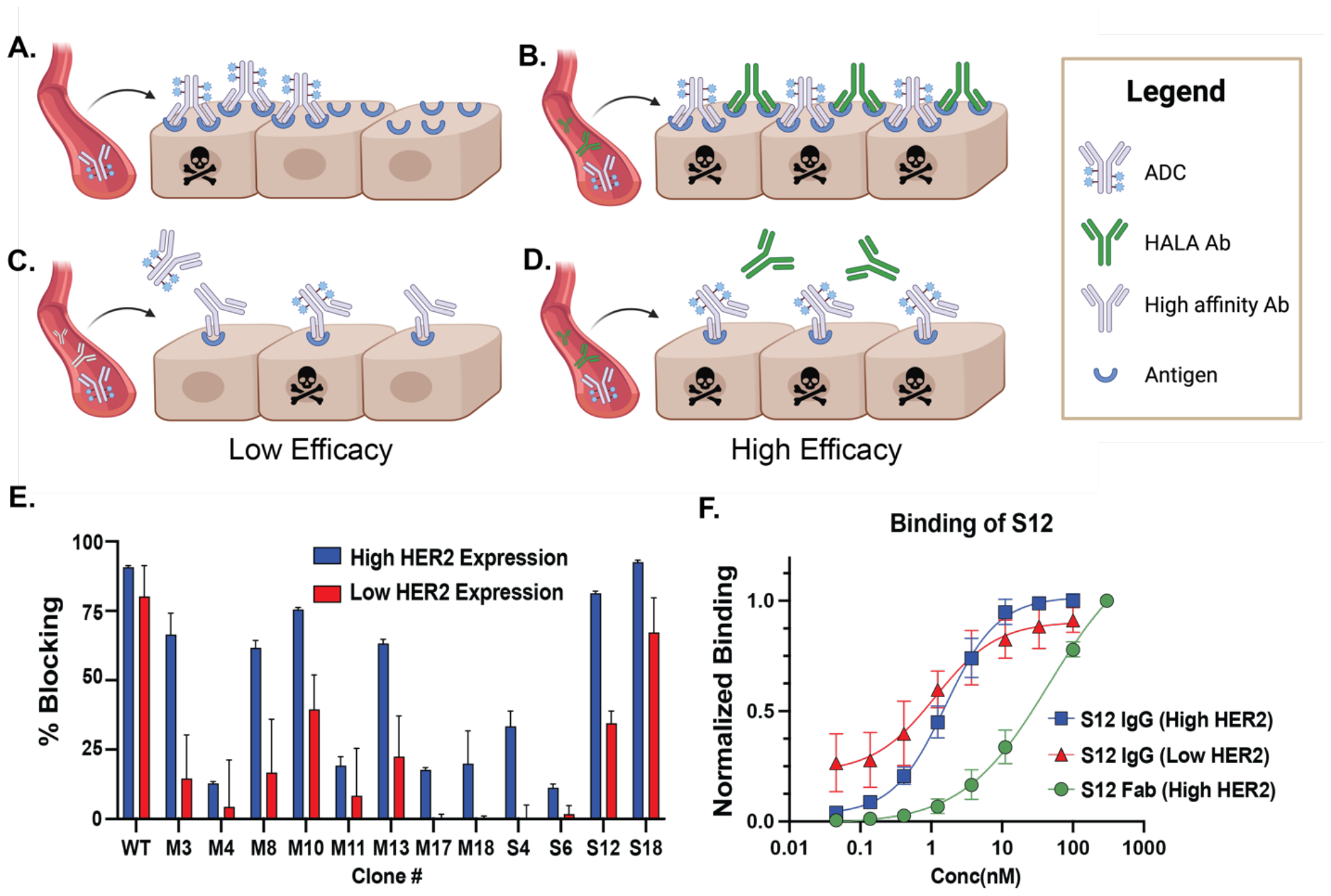
Schematic of HALA design approach to improve *in vivo* ADC distribution/efficacy and *in vitro* identification of engineered HALA antibody. **(A)** In high-expression tumors without a carrier dose, the ADC will reach a subset of cell layers beyond the blood vessel, limiting efficacy. **(B)** The addition of a carrier dose can partially block the binding sites, increasing tissue distribution and efficacy. **(C)** However, in a low-expression tumor, a high-affinity carrier dose competes with the ADC and prevents uptake into low-expression cells, reducing efficacy. **(B, D)** The use of a HALA carrier dose in **(B)** high-expression systems results in competition with the ADC due to high avidity interactions, but such competition is not observed in **(D)** low expression levels due to ‘auto-tuning’ the competition in situ. Therefore, HALA antibody intrinsically maximizes efficacy in both high and low expression systems. **(E)** Multiple HALA variants were tested for the optimum affinity on both high-expressing and low-expressing HER2 cell lines. The S12 variant maintained high competition on high expressing NCI-N87 cells but not on low-expressing MDA-MB-231 cells. **(F)** The HER2 binding of bivalent S12 (in the absence of an ADC) was high on both high expression HCC-1954 and low expressing MDA-MB-231 cell lines, but the Fab arm has a much lower intrinsic affinity.

ADC doses are typically limited by the amount of payload administered since the antibodies are often well tolerated,^28^ allowing high HALA doses relative to the ADC that can improve tissue penetration, leverage Fc effector functions^15^, and inhibit cancer signaling pathways (e.g., HER2). Given the effect of binding affinity on immune activity,^29–31^ we tested HALA antibodies with two FDA-approved ADCs, Kadcyla (T-DM1) and Enhertu (T-Dxd), in immunodeficient mouse models and a trastuzumab PBD ADC (T-PBD) in an immunocompetent model. Despite differences in payload potency, bystander effects, dosing, expression levels, and mouse models^32^,^33^, all ADCs demonstrated equal or greater efficacy when co-administered with the HALA antibody, highlighting the ability to enhance multiple mechanisms of action.

## RESULTS

### Generation and In vitro Binding Competition of HALA Antibodies

HALA antibodies were generated using a mutational library of trastuzumab on the surface of yeast, and lead clones were reformatted as IgGs. Binding competition between the HALA antibody panel and trastuzumab was measured in low (MDA-MB-231), medium (MDA-MB-453), and high (NCI-N87) HER2 expression cell lines (**Figure 1E**). A ratio of 8:1 for the competitor to ADC was selected based on the known ratio for efficient tissue penetration in vivo with a clinically relevant ADC dose of trastuzumab emtansine.^16,23^ From these results, the S12 clone was identified as the lead compound based on blocking in high expression systems (similar to the 90% blocking with trastuzumab as the carrier dose) with minimal blocking in low expression cells (only 35% binding despite an 8-fold higher concentration).

Binding affinity measurements were acquired with IgG and Fab fragments to measure bivalent and monovalent binding, respectively, on the cell surface (**Fig. 1F**). In a high expression system, the S12 HALA clone shows a K_D_ of 0.7 nM but only 46 nM when measured using the monovalent Fab arm, demonstrating high avidity. Surprisingly, the avidity remains high even on a low expression cell line. It is only when the higher affinity competitor is added that the HALA antibody is outcompeted on the cell surface, although this is consistent with theory.^23^

To further confirm that the HALA antibody was able to effectively block binding of T-DM1 ADC in an expression-dependent manner, cell viability assays were conducted by co-incubated trastuzumab or S12 with T-DM1(**Supplemental Figure S1**). S12 with T-DM1 resulted in a similar level of blocking as trastuzumab on a high expression cell line, while S12 resulting in less blocking and more cell killing on the lower expressing MDA-MB-453 cells as desired. Both cell lines had modest responses to trastuzumab alone (**Supplemental Figure S2**)

### HALA Antibody Improves 3D Tissue Penetration in Tumor Spheroids

To evaluate whether the binding competition on cell monolayers in vitro would translate to improved tissue penetration, we imaged the distribution of trastuzumab (as a surrogate for T-DM1 with no confounding cell killing) with or without 8:1 concentrations of HALA antibodies in high HER2 expression tumor spheroids (HCC-1954). Trastuzumab alone results in peripheral spheroid uptake due to poor tissue penetration, similar to T-DM1 (**Figure 2A**). In contrast, the addition of an 8:1 ratio of S12 HALA antibody partially blocks binding in the periphery, forcing deeper penetration of trastuzumab into the spheroid (**Figure 2B**). The M11 clone had even lower competition on the low expression MDA-MB-231 cell line compared to S12 (**Figure 1E**), but the competition was modest on the high expression cell line. When co-administered, trastuzumab remains bound at the periphery while the high concentration of M11 diffuses deeper into the tissue (**Figure 2C**), indicating the avidity effect is not sufficient to improve tissue penetration below a certain affinity.

**Figure 2.**
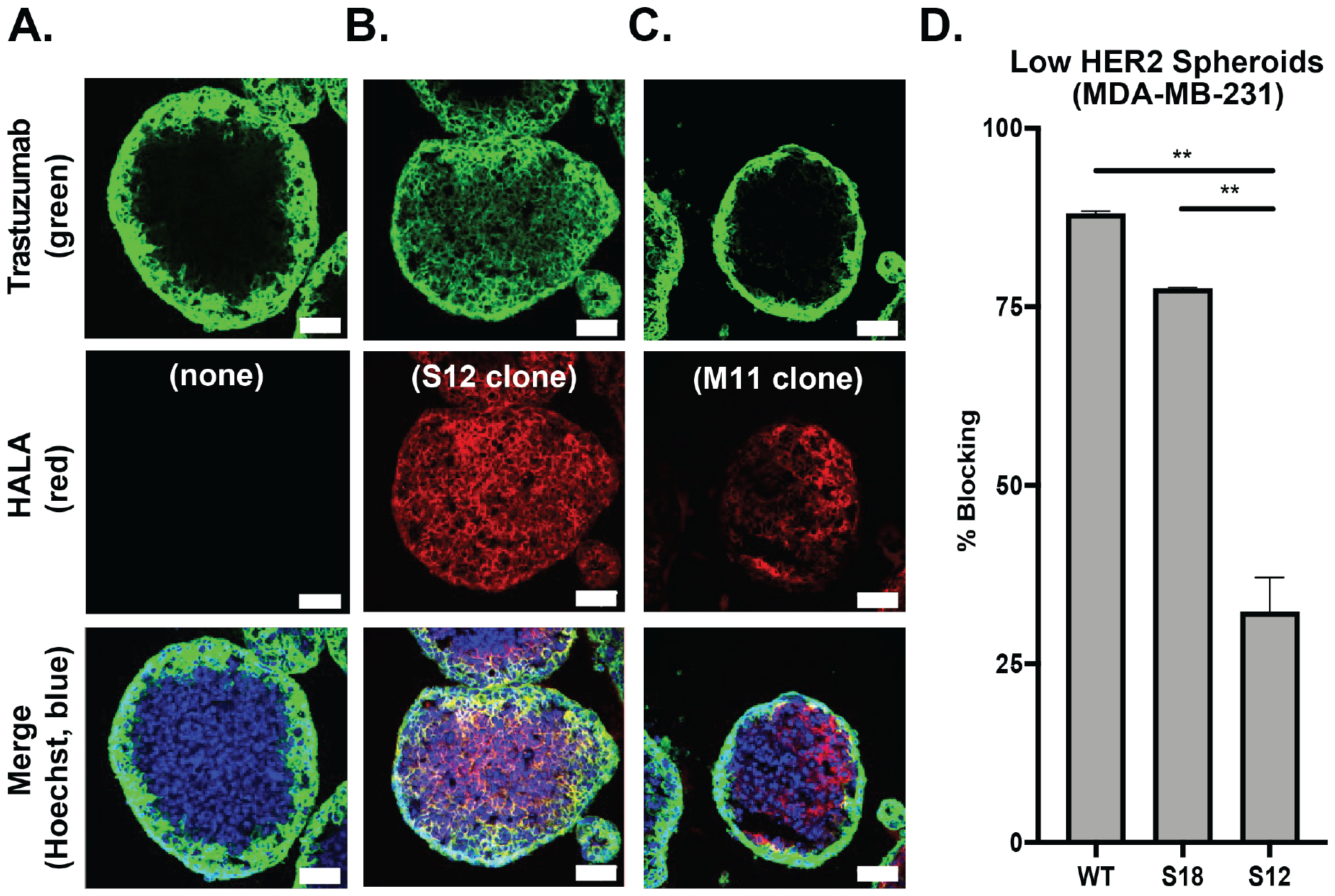
S12 HALA antibody ‘autotunes’ binding and distribution of trastuzumab in tumor spheroids. **(A)** trastuzumab (green; 50 nM) shows limited penetration into high HER2-expressing HCC-1954 spheroids (nuclei stained with Hoechst, blue). **(B)** The addition of the HALA antibody (8:1 S12:trastuzumab; red) to trastuzumab (50 nM, green) significantly enhanced trastuzumab tissue penetration. **(C)** In contrast, lower blocking HALA variants (e.g. M11) did not have sufficient affinity to compete with trastuzumab, resulting in only peripheral targeting of trastuzumab (green) despite M11 penetration to the spheroid center (red). **(D)** With low expression spheroids, clones with higher blocking than S12 (e.g. S18) reduce cellular uptake. Beneficially, S12 HALA shows low levels of competition with trastuzumab for low HER2-expression spheroids. Therefore, S12 has sufficient binding to improve tissue penetration in high expression spheroids **(B)** while maintaining low blocking in low expression spheroids **(D)**. Scale bar = 100 µm. Unpaired, two-sided t tests were used (** p<0.001).

Due to the low expression (and full tissue penetration) on the MDA-MB-231 cell line, MDA-MB-231 spheroids were digested into a single-cell suspension, and the average fluorescent signal was measured to quantify the competition between the carrier dose and ADC surrogate. Compared to several HALA test candidates, S12 showed the lowest binding competition with trastuzumab on this low expression cell line (**Figure 2D**). Therefore, the affinity of the HALA antibody must be balanced, since too high of an affinity (like the S18 clone or trastuzumab itself) will increase tissue penetration but block uptake in low expression cells, while too low of an affinity will not aid in tissue penetration (**Supplemental Figure S3**). S12 coincubation results in improved tissue penetration with slight gradients in trastuzumab uptake, indicating it has sufficient affinity to compete at high expression levels (**Supplemental Figure S4**). Together, the competition on the high expression cells to improve tissue penetration and low competition on the low expression cells made S12 the lead candidate for *in vivo* testing.

### HALA Antibody Improves *In Vivo* Tissue Penetration

We next measured the distribution of ADCs T-DM1 and T-Dxd *in vivo* with and without a HALA carrier dose to quantify the tissue penetration in xenograft tumors. Because tissue penetration is related to the dose/plasma concentration,^34^ we used the clinical doses of these ADCs to match the tissue penetration in the mouse model. Mice with high HER2 expression (NCI-N87) tumors were treated with T-DM1 or T-DXd, with and without a carrier dose, to determine how the HALA carrier dose impacted tissue penetration *in vivo*. For T-DM1, an 8:1 carrier dose (8-fold higher carrier antibody relative to the ADC, see methods section) of the S12 HALA antibody or trastuzumab improved the tissue penetration in vivo relative to T-DM1 alone (**Figure 3A-C**). The carrier doses (trastuzumab and the S12 HALA antibody) improved distribution as quantified with a Euclidean distance map (**Figure 3D**), with statistically significant differences in drug gradients (p < 0.005).

**Figure 3.**
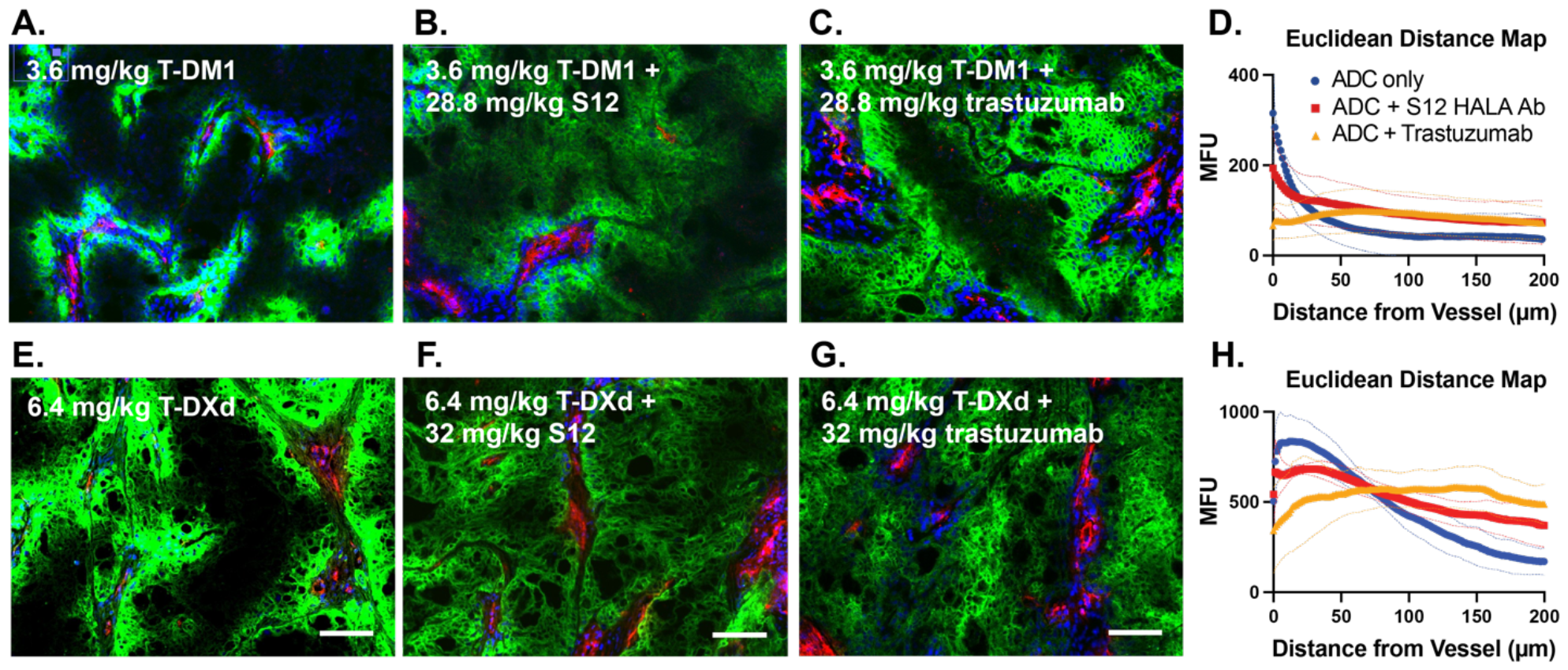
S12 HALA improves *in vivo* distribution of T-DM1 and T-DXd. **(A-C)** Representative histology images from mice (n=3) 24 hours post injection with **(A)** 3.6 mg/kg fluorescent T-DM1 (green) alone, **(B)** with an 8:1 ratio of S12 to T-DM1 or **(C)** with an 8:1 ratio of trastuzumab to T-DM1, showing improved ADC distribution with the addition of a carrier dose. Intravenous Hoechst (blue) labels nuclei around functional blood vessels, with all vessels stained with CD31 (red). **(D)** Euclidean distance mapping illustrates the mean fluorescence intensity versus distance from blood vessels from multiple images. **(E-G)** Representative histology images from mice (n=3) 72 hour post injection with **(E)** 6.4 mg/kg fluorescent T-DXd (green) alone, **(F)** with an 8:1 ratio of S12 to T-DXd or **(G)** with an 8:1 ratio of trastuzumab to T-DXd. Intravenous Hoechst (red) labels nuclei around functional blood vessels. **(H)** Euclidean distance mapping illustrates the mean fluorescence intensity versus distance from blood vessels from multiple images

T-Dxd uses a higher clinical dose of 6.4 mg/kg (in gastric cancer, 5.4 mg/kg in breast cancer), so the carrier dose ratios have to be reduced to maintain a similar total antibody dose for maximum tissue penetration. A 5:1 ratio was used to provide sufficient tissue penetration but prevent a reduction in total uptake (%ID/g) from oversaturating the tumor at an 8:1 ratio (**Supplemental Figure S5**). Similar to the carrier doses with T-DM1, the distribution was significantly improved when the ADC was co-administered with a 5:1 ratio of trastuzumab or S12 HALA antibody (**Figure 3E-H**). The differences in drug gradients for all 3 treatments were statistically significant (p < 0.0001).

### Co-administration of HALA Antibody with ADC Maximizes Tumor Growth Inhibition

After establishing that the S12 carrier dose can improve distribution in high expressing tumors in vivo but maintain minimal blocking in low expression tumors, we tested the impact of our HALA antibody on ADC efficacy. The goal was to improve the efficacy of the ADC in high expressing tumors while maintaining efficacy in moderate/low expressing tumors. Using high and moderate HER2 expression cell lines, T-DM1 was administered at its clinical dose of 3.6 mg/kg alone, with an 8:1 S12 carrier dose, or with an 8:1 trastuzumab carrier dose ratio. In the high expression NCI-N87 cell line, we observed the strongest tumor growth inhibition with the HALA carrier dose, which inhibited tumor growth more than a trastuzumab carrier dose or T-DM1 alone (**Figure 4A**). The trastuzumab or S12 carrier doses were better than T-DM1 alone by day 7 (p < 0.01) and maintained at day 14 (p < 0.0005). By day 21, the HALA carrier dose was performing better than the trastuzumab carrier dose (p = 0.011). In the low HER2 expression tumor model, MDA-MB-453, T-DM1 alone showed strong efficacy, as expected from better tissue penetration in lower expression tumors. The HALA carrier dose with ADC or ADC alone caused faster tumor regression than the trastuzumab plus ADC by day 7 (p < 0.02), (**Figure 4B**). The HALA carrier dose remained more effective at Day 14 but was no longer statistically significant by day 28 (p = 0.078), which could be due to the sensitivity of MDA-MB-453 cells to trastuzumab (**Supplemental Figure S2, Figure S6-S7**). In both cell lines, the HALA antibody on its own had some efficacy, which is promising from a clinical perspective (where a patient may benefit from multiple mechanisms of action including HER2 signaling blockade). Overall, the addition of S12 did not reduce efficacy, achieving the goal of the HALA approach.

**Figure 4.**
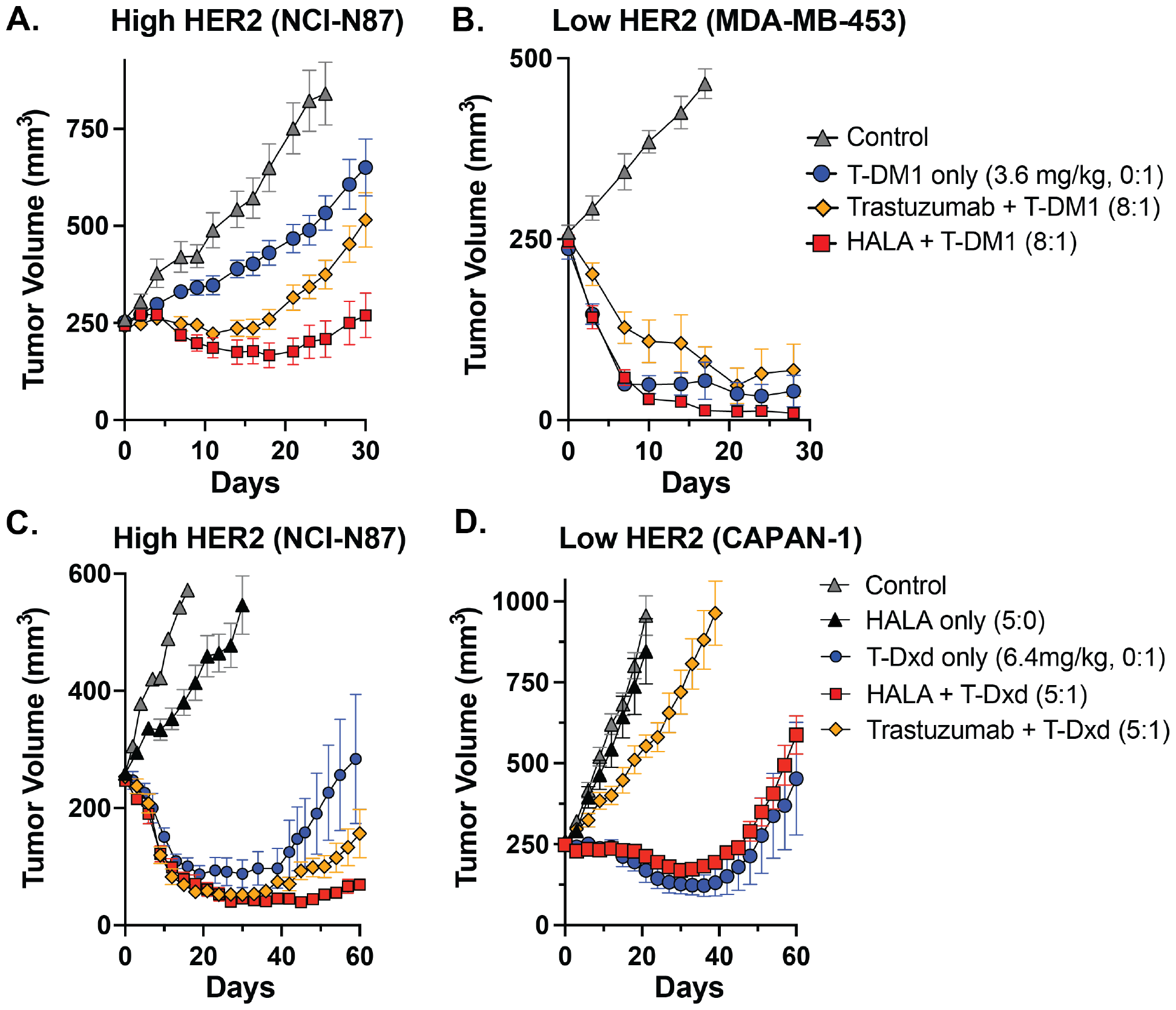
S12 HALA is uniquely effective at inhibiting tumor growth *in vivo* for both low- and high-HER2 expression tumors. **(A-B)** Tumor volume (mm^3^) versus time after treatment in **(A)** high HER2 expression (NCI-N87) cell line and **(B)** low expression (MDA-MB-453) cell line with T-DM1 monotherapy, T-DM1 with trastuzumab, or T-DM1 with S12 HALA antibody. **(C-D)** Tumor volume measurements were also made after T-DXd treatment in **(C)** NCI-N87 (high HER2 expression) or **(D)** Capan-1 (low HER2 expression) tumors alone, with T-Dxd and trastuzumab, or with T-DXd and S12 HALA. In **(A, C)**, the HALA antibody was able to improve ADC efficacy in high-expression tumors, similar to trastuzumab, but **(B, D)** the HALA administered with ADC maintained efficacy similar to ADC alone in moderate- and low-expressing tumors, unlike the trastuzumab carrier dose. Error bars indicate SEM.

Next, we tested the efficacy of T-DXd with a 5:1 ratio of S12 carrier dose based on improved distribution while maintaining total ADC uptake. T-DXd monotherapy showed higher baseline efficacy than T-DM1 monotherapy in the high expression NCI-N87 cell line due to higher ADC penetration at clinical doses and benefit of bystander payload penetration.^35,36^ The tumor cohorts started to differentiate at later times, with the HALA carrier dose showing smaller tumor sizes than the trastuzumab carrier (p = 0.041) and ADC alone (p = 0.054) at 8 weeks (**Figure 4C, S8**). In the low expression CAPAN-1 tumors (**Table S1**), the 5:1 trastuzumab carrier dose significantly lowered efficacy relative to ADC alone (p < 0.0001) due to competition at the low expression level, reducing ADC uptake and efficacy (**Figure 4D, S9**). However, S12 co-administered at a 5:1 ratio maintains similar efficacy to the ADC alone (p = 0.15 and 0.38 at 4 and 8 weeks), suggesting negligible competition for binding sites at low expression. Importantly, a dose of 32 mg/kg carrier dose alone shows no significant efficacy over negative controls, demonstrating that this xenograft model isolates the impact of payload delivery as a mechanism of action with the strongest benefit from the HALA antibody.

### Enhanced ADCC Activity with Fc-engineered HALA Antibodies

The xenograft models show how HALA antibodies improve efficacy in high expression tumors by increasing tissue penetration while preserving efficacy in low expression models. However, we wanted to engineer improved efficacy in low expression tumors as well as high expression. Therefore, we utilized another mechanism of action that can be leveraged to enhance killing on low expression cells – Fc effector function.

The high dosing and reduced affinity^29,37^ of a HALA antibody can benefit Fc-effector functions such as Antibody Dependent Cellular Cytotoxicity (ADCC) and Antibody Dependent Cellular Phagocytosis (ADCP). The co-administration of a HALA antibody with an ADC is expected to achieve a similar level of ADCC on targeted cells (i.e. perivascular cells) as ADC alone, but the higher dose from the HALA antibody will increase the tissue penetration and number of targeted cancer cells in vivo, thereby enabling greater ADCC in high expression tumors (**Figure 5A**). The HALA antibody would not impact ADCC with low expressing cancer cells due to a lack of binding/competition (**Figure 5B**). However, if the binding affinity of the HALA Fc-domain to Fc*γ* receptors is increased (with an enhanced Fc receptor binding domain, or eFc), the heterotypic avidity between HER2 on the cancer cell and Fc*γ*R on an immune cell can enable the HALA to outcompete the ADC in the immune synapse, generating greater ADCC than the ADC alone or ADC plus a wild type S12 or trastuzumab carrier dose (**Figure 5C**). Using an ADCC reporter cell line, WT antibody induces strong ADCC with or without the S12 carrier dose at a 3:1 ratio at high HER2 expression levels (**Figure 5D**). In a low HER2 expression cell line (∼100-fold fewer HER2 receptors/cell), inducing strong ADCC is more challenging with ∼7-fold lower induction with WT trastuzumab alone. However, the S12eFc carrier dose doubles the ADCC activity, demonstrating the ability of the HALA antibody to outcompete the ADC in the immune synapse (**Figure 5E**). This provides two mechanisms to enhance killing of low expression cells: in the absence of an immune cell, the ADC outcompetes the HALA antibody, maximizing payload delivery, while in the presence of an immune cell, the avidity from the Fc*γ* receptor and HER2 binding allow the HALA antibody to outcompete the ADC to maximize Fc-mediated killing.

**Figure 5.**
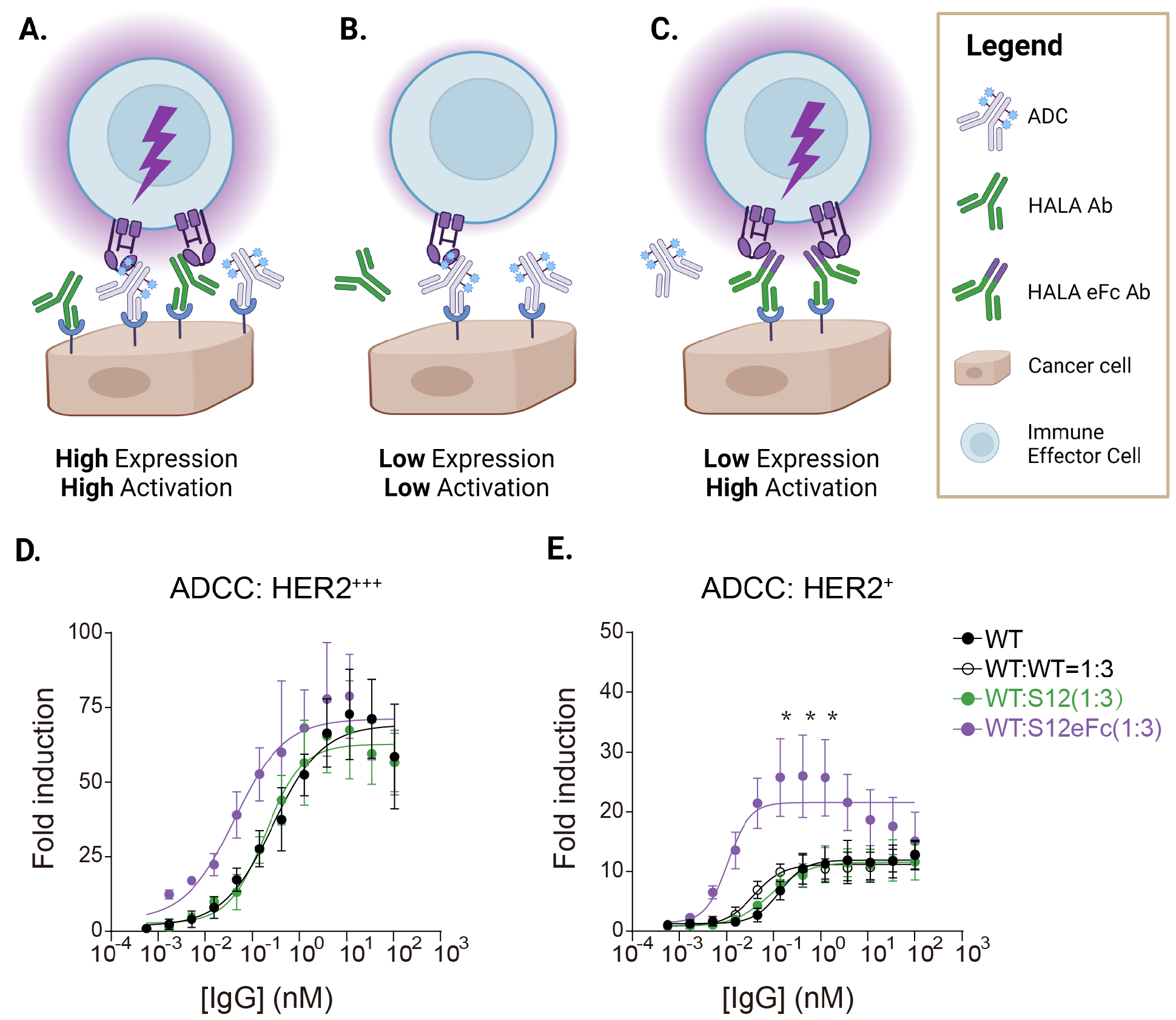
HALA antibody with an engineered Fc region improves the overall ADCC activity for both high- and low-HER2 expression cells. **(A)** The HALA antibody is expected to have minimal change in ADCC on high-expression cells that are already targeted, such as perivascular cells. However, it can improve overall ADCC at the tissue level by reaching more cells deeper in the tumor. **(B)** Lower HER2 expression cells naturally have lower ADCC, which is not impacted by the addition of a HALA antibody since the HALA antibody does not compete on low expression cells. **(C)** However, a higher-affinity Fc domain (referred to as enhanced Fc or eFc) on the HALA antibody can provide a boost in avidity between the Fc receptor and HER2 binding to enable binding in the context of immune cell synapses. Therefore, the higher Fc affinity can increase ADCC beyond a high-affinity antibody alone. **(D-E)** ADCC activity for trastuzumab alone or mixtures thereof was evaluated for **(D)** high HER2 (HCC-1954) and **(E)** low HER2 (MDA-MB-231) expression cells after incubation with trastuzumab (WT), a 3:1 ratio of S12 to WT, or a 3:1 ratio of S12eFc. In **(D)**, a similar maximum induction was observed (since there is no opportunity for improvement in tissue penetration on these cell monolayers), while in **(E)** the S12eFc:WT mixture shows a large increase in ADCC relative to WT alone (p < 0.0005), which was only ∼3-fold lower than induction with the high expression cell line despite a ∼100-fold lower HER2 expression.

### Co-administered HALA Antibodies Induce Strong Responses in a Syngeneic Mouse Model

To test the ability of the enhanced Fc HALA antibodies to improve efficacy in an immunocompetent setting, we used E0771 cells transfected with human HER2 (hHER2) and injected them into hHER2 transgenic mice. Mouse cells are less sensitive to DM1 or DXd payloads, requiring large doses for efficacy in syngeneic mouse models that can obscure the need for a carrier dose.^38,39^ Therefore, we used PBD as the payload for the syngeneic model given the high potency in both mouse and human cells. A 1 mg/kg of Trastuzumab-PBD (T-PBD, DAR 2) dose was selected to provide a moderate response for assessing changes in efficacy. Mice treated with 1 mg/kg of T-PBD showed a 60% overall response rate including 4/10 complete responses and 2/10 partial responses (**Figure 6**). To test the impact of Fc-effector function in this model, the Fc-effector function was eliminated by mutating the Fc-domain on the ADC (**Supplemental Figure S10**). Mice treated with 1.1 mg/kg of LALAPG-T-PBD (to match the same total payload dose) showed growth delay but only a 30% overall response rate and 1/10 mice with a complete response. In contrast, mice treated with 1 mg/kg T-PBD (containing a wild type Fc domain) co-administered with 3 mg/kg of S12eFc HALA antibody had the highest overall response rate with 82% including 4/11 complete responses and 5/11 partial responses.

**Figure 6.**
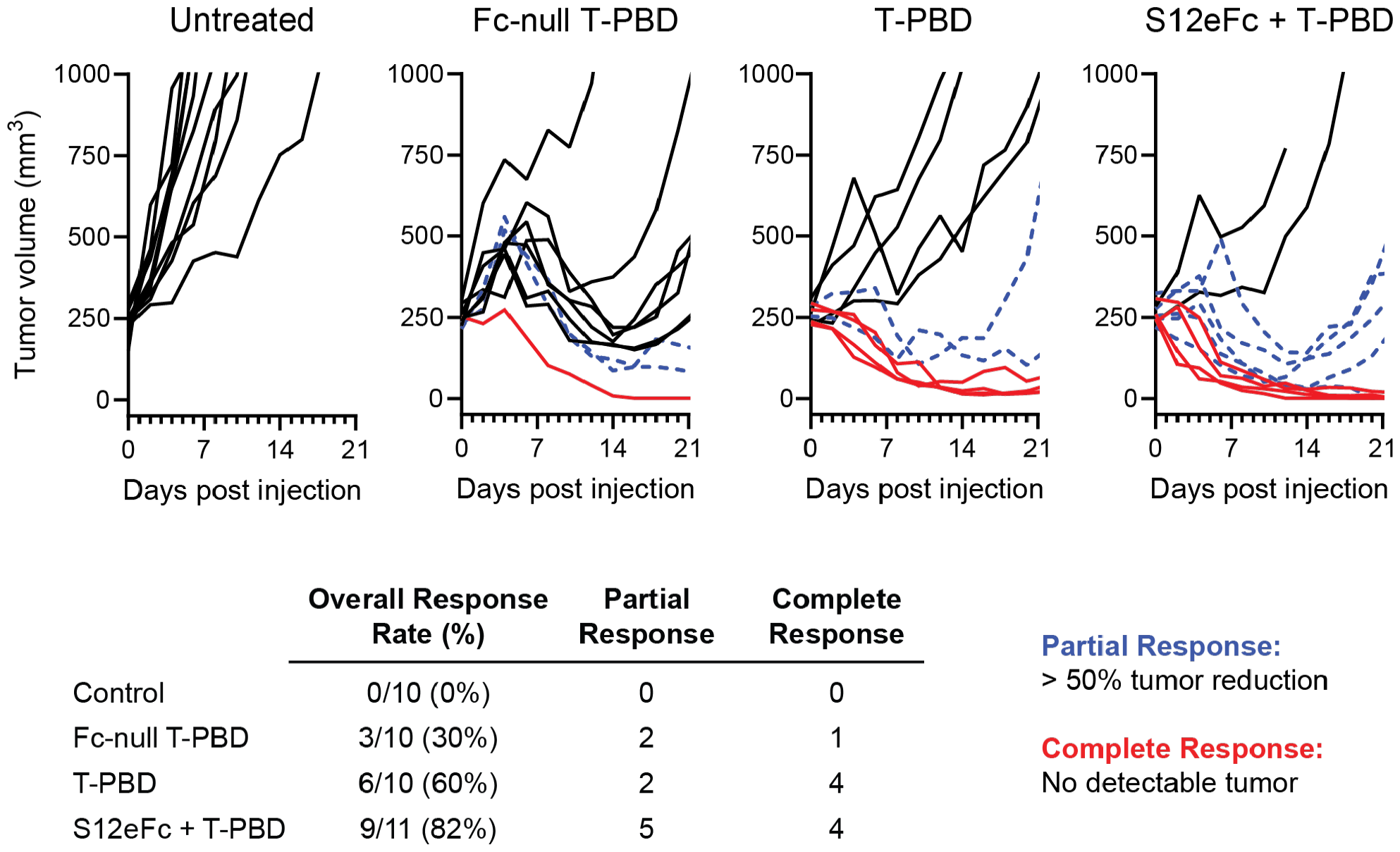
A mixture of ADC and S12 HALA antibody with an enhanced Fc domain shows strongest tumor growth inhibition in an *in vivo* syngeneic tumor model. Using a syngeneic mouse model, the impact of an enhanced Fc domain that has increased Fc-effector function was tested *in vivo*. Trastuzumab was conjugated to the DNA alkylating payload PBD (T-PBD) to treat the syngeneic model since T-DM1 and T-DXd have minimal potency in the mouse E0771 cell line. Human HER2-transfected E0771 cells were orthotopically implanted in the mammary fat pads of human HER2 transgenic mice. When tumors reached 250 mm^3^, mice were treated with a single injection of either 1.1 mg/kg of an Fc-null (LALA-PG mutated) trastuzumab-PBD ADC (DAR 2), 1 mg/kg of trastuzumab-PBD (containing a WT Fc domain), or 1 mg/kg of trastuzumab-PBD plus 3 mg/kg of an enhanced Fc HALA antibody. The Fc-null T-PBD showed a 30% response rate and one complete response, while the T-PBD had much higher efficacy with a 60% response rate and four complete responses. However, the most effective treatment was the T-PBD co-administered with the enhanced Fc HALA antibody, resulting in an 82% response rate including four complete responses.

## Discussion

Tumor penetration is a persistent challenge in ADC development and is critical to achieving clinical responses in solid tumors. Poor tissue penetration has sparked multiple approaches such as bystander payloads,^40^ transient anti-drug binding,^13^ smaller format protein-drug conjugates,^10^ lower potency payloads to allow a larger antibody dose,^41^ and co-administration of antibody with an ADC to improve efficacy.^7,8,16–18^ A major hurdle with any of these approaches is the variable expression found with many targets. If target expression were homogeneous, a patient with high expression could receive an antibody carrier dose, while a patient with low expression could receive the ADC alone. However, this is not possible when expression levels differ between tumors/metastases or within the same tumor. Here, we designed a novel HALA carrier dose to co-administer with an ADC that ‘auto-tunes’ the level of ADC binding to match the amount of payload needed for efficient cell death (**Figure 1**). We tested the tissue penetration and efficacy with two clinically-approved ADCs to modulate cellular targeting to the expression level and better leverage multiple mechanisms of action. It was important to balance the HALA dose and affinity for T-Dxd and T-DM1 since a low HALA dose would not enhance penetration, but a saturating dose, where ADC uptake becomes limited by available binding sites,^42^ would decrease ADC tumor delivery (**Supplemental Figure S5**). Overall, HALA co-administration resulted in the largest and most consistent efficacy relative to ADC with or without a carrier dose. This approach can be applied to any ADC with high affinity and variable target expression provided the ADC is sufficiently potent to kill low/moderate expression cells. Therefore, the approach can broaden the use of carrier doses to clinically relevant scenarios where receptor expression varies within the same tumor or between metastases.^21^

There has been great interest in engineering ADCs for low to moderately expressed targets for ADC development,^43^ but even moderately expressing tumors can show significant differences in efficacy based on dosing and tissue penetration.^44^ Additionally, there has been a shift in ADC design to consider multiple mechanisms of action, such as payload delivery, immunogenic cell death (ICD), antibody signaling blockade, and Fc effector functions that could act synergistically to bolster efficacy.^45^ HALA antibody co-administration has the potential to improve all these mechanisms by increasing payload delivery/ICD, blocking receptor signaling (e.g. HER2), and contributing to ADCC (as seen with trastuzumab^14^). With the two approved HER2 ADCs, T-DXd monotherapy shows stronger efficacy than T-DM1, which benefits from higher dosing and payload bystander effects^46^. Bystander effects provide some improvements against the binding site barrier^20,47^, but tissue penetration has been shown to be important even with bystander payloads.^17,36^ Therefore, increased penetration in the high expression model further improved T-DXd efficacy.

The increased efficacy of an enhanced Fc HALA antibody (utilizing the same Fc mutations as margetuximab^48^) was demonstrated in a syngeneic mouse model of breast cancer, highlighting the ability of HALA antibodies to further leverage the immune response. The HALA antibody is therefore capable of maximizing HER2 signaling blockade (as seen with efficacy in the moderate expressing MDA-MB-453 model), increasing payload delivery and ICD/cell death, and increasing Fc-effector function as seen in the syngeneic mouse model.

Overall, we demonstrate how HALA antibodies can improve ADC treatment efficacy through multiple mechanisms of action at both high and low expression levels. We achieved favorable ADC distribution and cellular uptake across different expression levels using HALA carrier doses that ‘auto-tune’ the delivery at the cellular level, addressing a key limitation and expanding the use of carrier doses to a greater number of clinically relevant scenarios. ADCs have been increasingly developed as immunotherapies rather than a payload delivery mechanism, and this approach enhances the contribution of immune effects to provide stronger and more durable ADC responses.

## Acknowledgments

This work was supported by the Department of Defense Breast Cancer Research Program Grant #BC200857 (GMT, PMT, and JJL) and the Albert M. Mattocks Chair fund (PMT). Research reported in this publication was also supported by the National Cancer Institutes of Health under Award Number P30 CA046592 by the use of the following Cancer Center Shared Resource(s): Flow Cytometry, Tissue and Molecular Pathology.

## MATERIALS AND METHODS

### Sorting of high avidity, low affinity (HALA) trastuzumab mutants

The trastuzumab antibody fragment was displayed on the surface of yeast along with a library of mutant sequences with several mutations per antibody. For sequence generation, a heavy chain complementarity determining region (HCDR) focused library for trastuzumab was made in single chain Fab (scFab) format as described previously^49^ to isolate trastuzumab mimics with high avidity, low affinity. We targeted sites 53 and 56 in HCDR2 and 96, 97, 98, 99, 100, 100A and 100B in HCDR3. The library DNA for scFab was assembled by PCR, transformed with modified pCTCON2 plasmid in yeast strain EBY100 by homologous recombination as described.^50^ The designed library size was ∼10^7^ and we achieved >10^8^ transformants.

To enrich the library for antibodies with affinities in the desired range, we selected clones that were able to bind a bivalent antigen (HER2-Fc fusion) at a similar level as wild type trastuzumab but had reduced binding to monovalent HER2. The library was sorted against bivalent HER2-Fc antigen immobilized on dyna beads by magnetic activated cell sorting (MACS) in round 1 and 2. In round 1, 10^9^ yeast cells displaying library were incubated with 10^7^ beads coated with bivalent HER2-Fc for 3 hours at room temperature in 1% milk or 1% serum in PBS+0.1% BSA (PBSB). Post incubation the beads and unbound cells were separated by magnetic separation.^50^ The beads were washed once with ice-cold PBSB and then re-suspended in 50 mL SDCAA growth media and grown overnight at 30 °C. In round 2, MACS was performed as described for round 1 but starting with 10^7^ yeast cells and 10^7^ beads.

In round 3, the libraries were sorted by florescence activated cell sorting (FACS) against monovalent HER2 fused to Fc domain of human IgG1 (HER2-mFc). 10^7^ yeast cells were incubated with 100 nM of HER2-mFc, 1/1000x dilution of mouse anti-myc antibody and incubated in either 1% milk or 1% mouse serum in PBSB for 3 hours at room temperature. The cells were then centrifuged, washed once with ice-cold PBSB followed by incubation with secondary reagents; 1/100x dilution of goat anti-mouse Alexa Fluor 488 and 1/300x dilution of goat anti-human Fc Alexa Fluor 647 on ice for 4 mins. Post incubation, the cells were washed once with ice-cold PBSB and sorted on Beckman Coulter Astrios sorter. Approximately 40 individual clones were tested on yeast (**Supplemental Figure S11**), and 12 clones were selected for characterization of binding competition on mammalian cells.

### Tuning HALA Antibody Binding Affinity

Reformatted IgG was serially diluted in PBS with 0.5% BSA starting from 300nM and added to 96 well u-bottom plates at 100µl/well. HCC1954 and MDAMB231 cells were trypsinized, resuspended in RPMI1640, spun down, washed in PBS-BSA, and added at 50,000 cells/well in 100µl. Plates were incubated with the antibody on ice for 4 hours. After incubation, cells were washed with PBS-BSA, labeled with Alexa anti-human Fc (Jackson Immuno 109-605-190) at a 300x dilution for 4 min on ice, washed in PBS-BSA, and run on flow cytometry (Bio-Rad Ze5). Data was plotted using Prism (GraphPad).

Fab fragments were concentrated to ∼2mg/ml in 100 µl PBS and labeled with 2 µl of 10 mg/ml Alexa-647 (Fisher Scientific A37573), reacted for 1 hr, and purified with 10% P6 Biogel (Bio-Rad 1504130) in PBS loaded to a costar spin-x tube (Corning 07-200-387). The protein concentration and degree of labeling were then measured on the nanodrop. The affinity of the fragments was measured as above.

### Cell culture and animals

NCI-N87, HCC1954, MDA-MB-453, and MDA-MB-231 cell lines were obtained from ATCC and maintained according to ATCC guidelines. Cells were passaged 2-3 times per week up to passage 50 using RPMI 1640 media (NCI-N87, HCC1954) or DMEM (MDA-MB-453, MDA-MB-231) each supplemented with 10% (v/v) FBS, 50 U/mL penicillin and 50 µg/mL streptomycin. Cells were incubated at 37 ºC in 5% CO_2_. 6 to 8 week old female nude mice were obtained from Jackson Labs and all studies were carried out according to and with approval of University of Michigan IACUC and AALAC guidelines.

### In vitro HALA and trastuzumab binding competition

Trastuzumab was fluorescently labeled with AlexaFluor 647 with NHS-ester chemistry as described previously.^51^ HCC-1954, NCI-N87, MDA-MB-453, and MDA-MB-231 cells were seeded at 300,000 cells/well in a 24 well plate and allowed to adhere overnight. The following day, media was replaced with 500 µL containing 5 nM fluorescent trastuzumab and 40 nM unlabeled HALA antibody, 5 nM trastuzumab and 40 nM unlabeled trastuzumab, or 5 nM fluorescent trastuzumab only for an 8-hour incubation at 37 ºC. (These resulted in an 8:1 ratio, where the carrier dose/concentration is always listed first with the ADC or ADC surrogate antibody listed second for all experiments.) Cells were then washed twice with PBS and detached from wells using Trypsin. Cells were washed again in PBS supplemented with 0.5% bovine serum albumin (BSA), then analyzed on an Attune NxT flow cytometer. Each condition was completed in duplicate and biological replicates were performed in triplicate. The percent blocking was calculated by subtracting autofluorescence from negative samples and was calculated by: (signal from trastuzumab only – signal with blocking Ab)/signal from trastuzumab only. Data was plotted using Prism (GraphPad).

### 3D tissue penetration in tumor organoids

Tumor organoids were grown as described previously.^10,52^ Briefly, HCC1954 or MDA-MB-231 cells were seeded at 3,000 cells/microwell and maintained by replacing media every other day. A week after seeding, spheroids were incubated with a mixture of 50 nM trastuzumab labeled with AlexaFluor 647 (AF647) and 400 nM M10, S12, or S18 antibodies labeled with AlexaFluor AF750 (AF750) for an 8:1 ratio of carrier dose antibody to trastuzumab (ADC surrogate). HCC1954 spheroids were harvested and flash frozen in OCT, then sectioned on a cryostat. Spheroids were stained for 1 minute with Hoechst 33342 to mark all cell nuclei and imaging was performed on an Olympus FV1200 confocal microscope using the 405 nm, 635 nm, and 750 nm lasers. Image analysis was performed in Fiji and Euclidean distance mapping of spheroids was completed in MATLAB.

Low HER2 expression MDA-MB-231 spheroids were digested into a single cell suspension by incubating n=20 spheroids of each condition with 0.05% trypsin-EDTA until spheroids were broken into single cells. The single cells were washed twice and passed through a 40 µm filter to remove cells clumps. Cells were analyzed using an Attune NxT flow cytometer to determine the amount of blocking based on reduction of trastuzumab-AF647 signal. Data shown is the result of three biological replicates consisting of n=20 spheroids for each independent experiment.

### In vivo distribution studies

Tumor-bearing nude mice were injected with fluorescent ADC and HALA antibody when tumors reached 250 – 300 mm^3^. Mice were sacrificed 24 hours after injection and tumors were resected for single-cell digests or embedded in OCT and flash frozen in isopentane for histology. Single-cell suspensions were prepared using Tumor Dissociation Kits (Miltenyi Biotec), filtered twice through 40 µm filters to remove clumps, and analyzed using an Attune NxT flow cytometer. The total payload in each bin was calculated by the following equation: (% of cells in each bin)*(median signal per bin)/(fluorescent signal per ADC) to create a weighted average of ADC delivery. The bins were determined based on correlations between fluorescent signal and IC_50_, where the ‘low’ bin corresponds to concentrations lower than the IC_50_, and the threshold for the ‘high’ bin is 10x the IC_50_.

Frozen tumors for histology were sectioned using a cryostat into 12 µm slices. Sections were stained with CD31-AF555 for 30 minutes in PBS-BSA, then washed with PBS before imaging. Microscopy was performed using a 20x objective on an Olympus FV1200 microscope using 405 nm, 543 nm, and 635 nm lasers and images were analyzed in ImageJ.

### In vivo efficacy studies

For the efficacy study with T-DM1, nu/nu mice (Jackson Labs) were injected with 5 million cells of either MDA-MB-453 (moderate HER2 expression) or NCI-N87 (high HER2 expression) into the left flank with 50% v/v Matrigel. Tumor volume was measured using calipers and the formula length^2^ x width/2 where length is the shorter dimension of the tumor. Once tumors reached a volume of approximately 250 mm^3^, tumor bearing mice were administered one of the following treatments via tail vein injection: 1) PBS vehicle control, 2) 3.6 mg/kg T-DM1, 3) 3.6 mg/kg T-DM1 and 28.8 mg/kg S12 (8:1 ratio of carrier antibody to ADC), 4) 3.6 mg/kg T-DM1 and 28.8 mg/kg Trastuzumab, or 5) 28.8 mg/kg S12. Tumors were monitored twice per week for MDA-MB-453 tumors and three times per week for NCI-N87 tumors until the humane endpoint. Data were plotted using PRISM (GraphPad).

For the efficacy study with T-DXd, 5 million cells of CAPAN-1 (moderate HER2 expression) or NCI-N87 were injected to each nude mouse, and tumor volumes was measured as previously described. Once tumors reached a volume of approximately 250 mm^3^, CAPAN-1 and NCI-N87 tumor bearing mice were administered one of the following treatments via tail vein injection: 1) PBS vehicle control, 2) 6.4 mg/kg T-Dxd, 3) 6.4 mg/kg T-Dxd and 32 mg/kg S12 (1:5 ratio), 4) 6.4 mg/kg T-Dxd and 32 mg/kg Trastuzumab, or 5) 32 mg/kg S12 with n=8 mice for each group. Tumors were monitored once every three days until the humane endpoint. Data were plotted using PRISM (GraphPad).

### ADCC Assays

ADCC cells (Invivogen, jktl-nfat-cd16) were cultured in IMDM media (Fisher 12440053) with 10% HI FBS, 1% P/S and 100 µg/ml Normocin (Invivogen ant-nr-1). 10 µg/ml of Blasticidin (Invivogen ant-bl-05) and 100 µg/ml of Zeocin (Invivogen ant-zn-05) were added to the growth media every passage. HCC-1954 or MDA-MB-231 cells were seeded to 96 well Tissue culture-treated flat-bottom plates as 50,000 cells/well and incubated at 37°C and 5% CO2 overnight. Antibodies were diluted as a 3-fold series starting from 100 nM, blocking antibodies were added as the required ratio if needed. After adding 110 µl of diluted antibody to each well, cells were incubated at 37°C and 5% CO_2_ for 1 hour. ADCC reporter cells were suspended in test medium (IMDM media with 10% HI FBS, 1% P/S), added to the wells (0.2 million cells in 90 µl) and incubated at 37°C and 5% CO_2_ for 6 hours. After incubation, 20 µl per well was placed in a white assay plate (Fisher 720091), 50 µl of QUANTI-Luc™ (Invivogen REPQLC2) was added per well, and luminescence was measured immediately.

### Syngeneic Mouse Model and Treatment

Efficacy studies were also conducted in an immunocompetent syngeneic mouse model. C57BL/6 hHER2 transgenic mice were obtained from Jackson Laboratory, and the breeding colony was maintained by the Unit for Laboratory Animal Medicine (ULAM) at the University of Michigan. 6-8 weeks old hHER2 transgenic mice were injected with 5 × 10^6^ cells of human HER2 (hHER2) transfected E0771 (E0771-hHER2) into the 4^th^ mammary fat pad with 50% v/v Matrigel. Tumor volume was measured as mentioned above. Once the tumor volume reached approximately 250 mm^3^, E0771-hHER2 tumor bearing mice were administered one of the following treatments via tail vein injection: 1) no treatment (n=7), 2) 1 mg/kg T-PBD (n=10), 3) 1.1 mg/kg LALAPG-T-PBD (to match the payload concentration) (n=10), 4) 1 mg/kg T-PBD and 3 mg/kg eFc-HALA S12 antibody (n=11). Tumors were measured every other day until the humane endpoint. Tumors that were not measurable and did not regrow by day 60 were considered complete responders. 5 × 10^6^ cells of E0771-hHER2 and E0771 wild type were subcutaneously injected into either the 2^nd^ or the contralateral 4^th^ mammary fat pad. No additional treatment was given. The tumors were measured every other day until the humane endpoint. Data were plotted using PRISM (GraphPad).

