## Supplemental Figures and Tables for "*In vivo* Auto-tuning of Antibody-Drug Conjugate Delivery for Effective Immunotherapy using High-Avidity, Low-Affinity Antibodies"

### Supplementary Information

### Methods

#### Cell viability assays

Cell viability assays were performed using NCI-N87 cells and MDA-MB-453 cells seeded at 6,000 cells/well in a black-wall, clear bottom 96 well plate and allowed to adhere overnight. Media was replaced daily with serial dilutions of T-DM1, S12, or Trastuzumab, or a combination of increasing T-DM1 concentration and carrier antibody (S12 or trastuzumab) maintaining a total antibody concentration of 10 nM. Media with antibody or ADC solutions was replaced daily for 6 days. On the final day, all media was removed and replaced with 10X PrestoBlue viability reagent diluted in complete media. Cells were incubated with PrestoBlue for two hours before scanning the plate on a BioTek Plate reader with 560/590nm excitation/emission. IC50 curves were fit in Prism (GraphPad).

Supplementary Figures

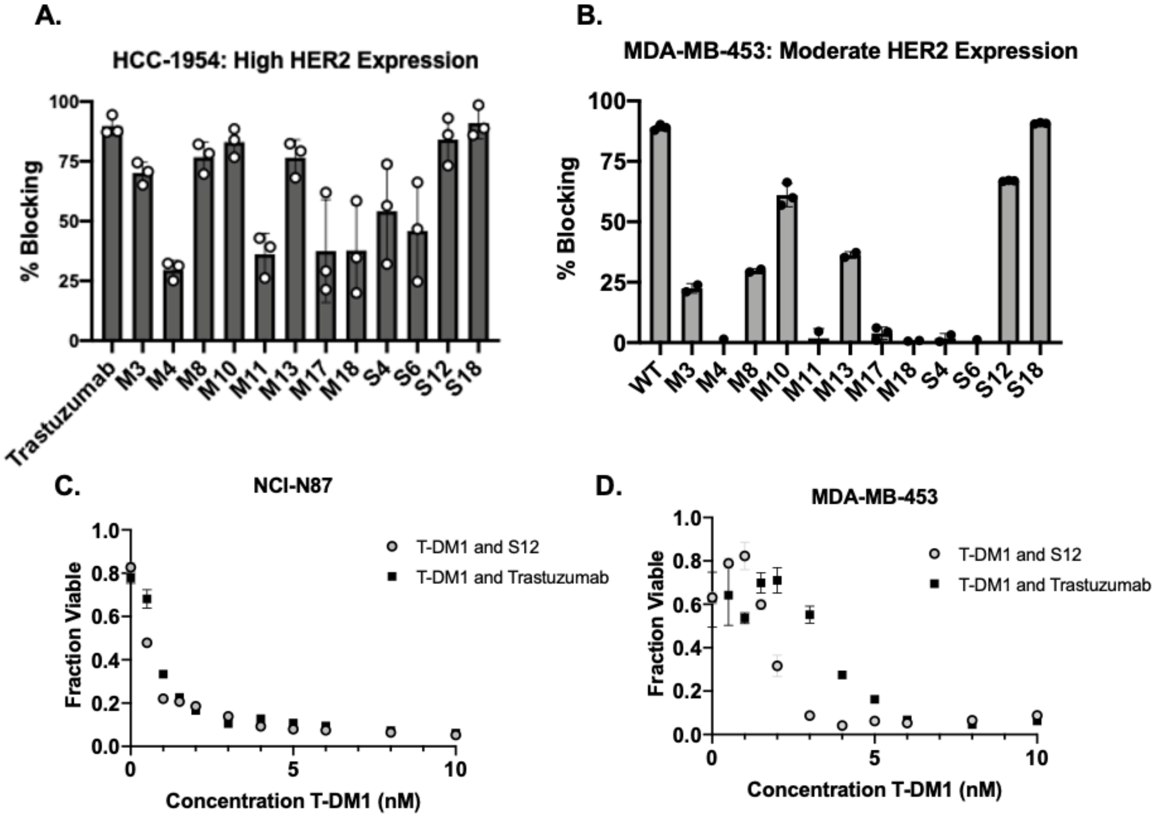


**Figure S1: Binding competition and Cell Viability of Antibody/ADC Combination.** Binding competition on (A) high HER2 expression HCC-1954 and (B) moderate HER2 expression MDA-MB-453 cells. Fraction of cells viable after incubation with varying concentrations of T-DM1 and carrier doses for a total of 10 nM antibody in high (C) and moderate (D) HER2 expression cell lines.

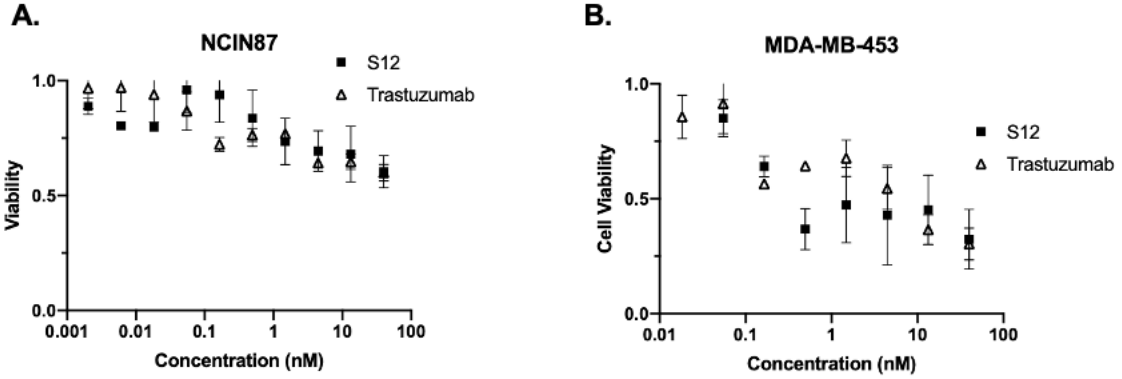


**Figure S2: Cell Viability Assays with Antibody Monotherapy.** Cellular viability assays with trastuzumab and S12 in (A) NCI-N87 and (B) MDA-MB-453 cell lines.

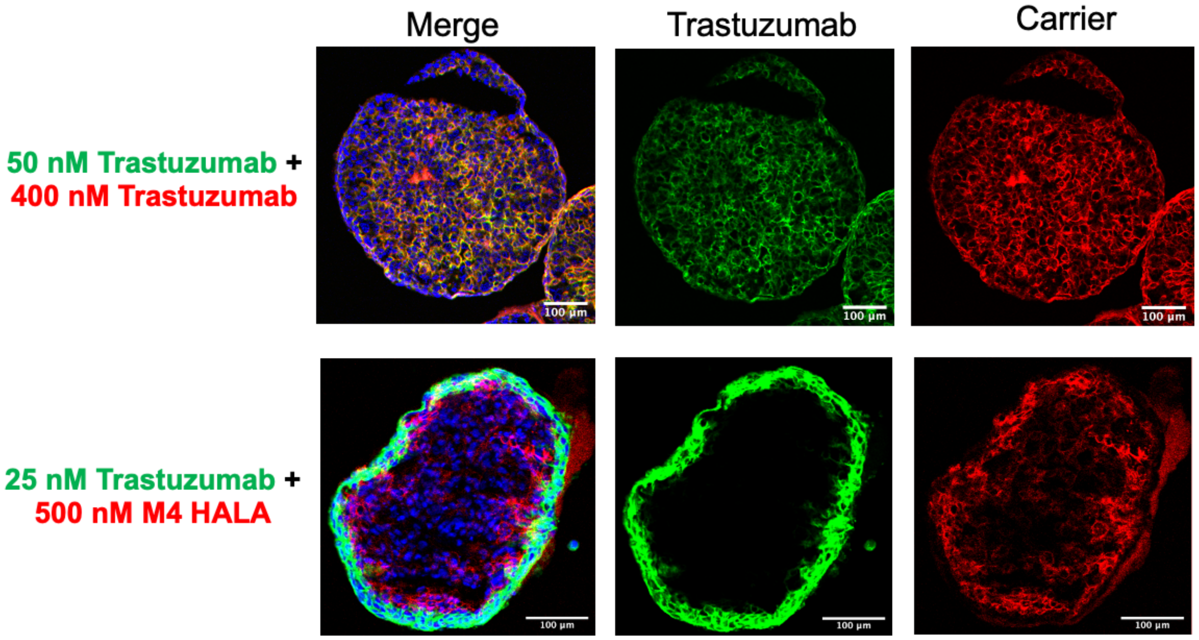


**Figure S3: Spheroid Penetration of Trastuzumab-AF647 Increases with Co-administered Trastuzumab-AF750 But Not M4 HALA-AF750**. Trastuzumab tested at an 8:1 ratio in HCC1954 spheroids (top row). Additional HALA clone, M4, was tested in spheroids at a 20:1 ratio (bottom row). Spheroids incubated with HALA-AF750 (red) and trastuzumab-AF647 (green) have cell nuclei are stained with Hoechst 33342 (blue).

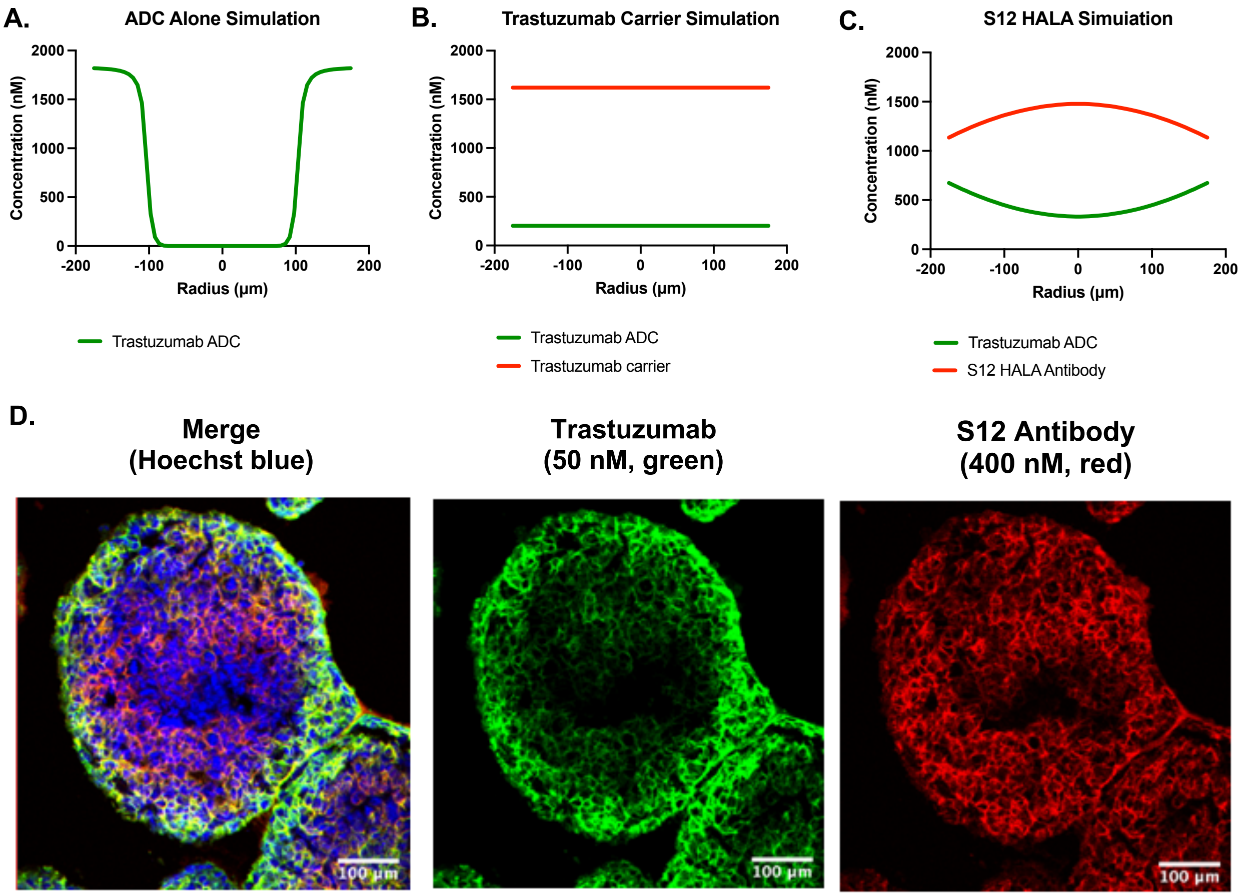


**Figure S4: Spheroid Simulations and S12 HALA Antibody Spheroid Imaging**. (A) Simulations of a trastuzumab antibody or ADC alone (50 nM in HCC1954 spheroids) results in peripheral binding in the spheroids due to rapid binding relative to diffusion. (B) The addition of 8:1 trastuzumab antibody to the 50 nM ADC results in more uniform ADC distribution. However, this would strongly compete with lower expression spheroids. (C) The addition of the S12 HALA antibody at an 8:1 ratio to 50 nM ADC results in a more uniform distribution of ADC with some curvature in the concentration profile due to incomplete competition of the HALA antibody for binding. Simulations used the same equations and code as Evans and Thurber, 2022, Sci Reports (D) HCC1954 tumor spheroids incubated for 24 hr with an 8:1 ratio of S12 HALA antibody (red) to 50 nM trastuzumab (green) show a similar slight gradient in binding while ensuring efficient tissue penetration of trastuzumab to the spheroid center.

.

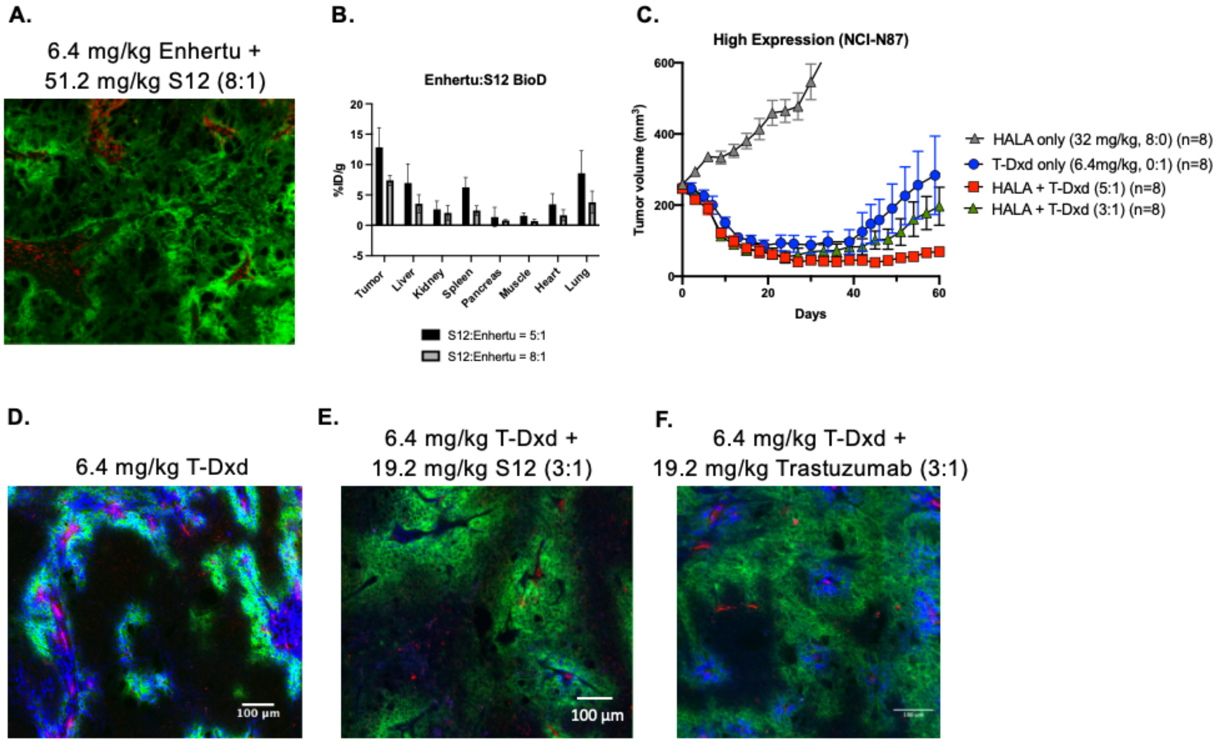


**Figure S5: Impact of HALA Dose on Tumor Uptake and Distribution**. Images of T-DXd and S12 at an 8:1 ratio (A) show high tumor tissue distribution, but biodistribution (B) suggests oversaturated tumors due to the decrease in %ID/g in the tumor at the 8:1 ratio compared to 5:1. (C) Tumor growth at 3:1 ratios of S12 to T-Dxd (green) shows decreased efficacy compared to the 5:1 ratio (T-DXd and 5:1 S12 data are duplicated from Figure 3 for comparison). (D) After 24 hrs, a 6.4 mg/kg dose of T-DXd alone shows poor tissue penetration. (E) The coadministration of 19.2 mg/kg S12 HALA with T-DXd (3:1) slightly improves penetration, but not as much as the coadministration of 19.2 mg/kg trastuzumab with T-DXd (3:1) (F). This indicates the 3:1 ratio of the S12 HALA antibody does not sufficiently increase tissue penetration to improve efficacy.

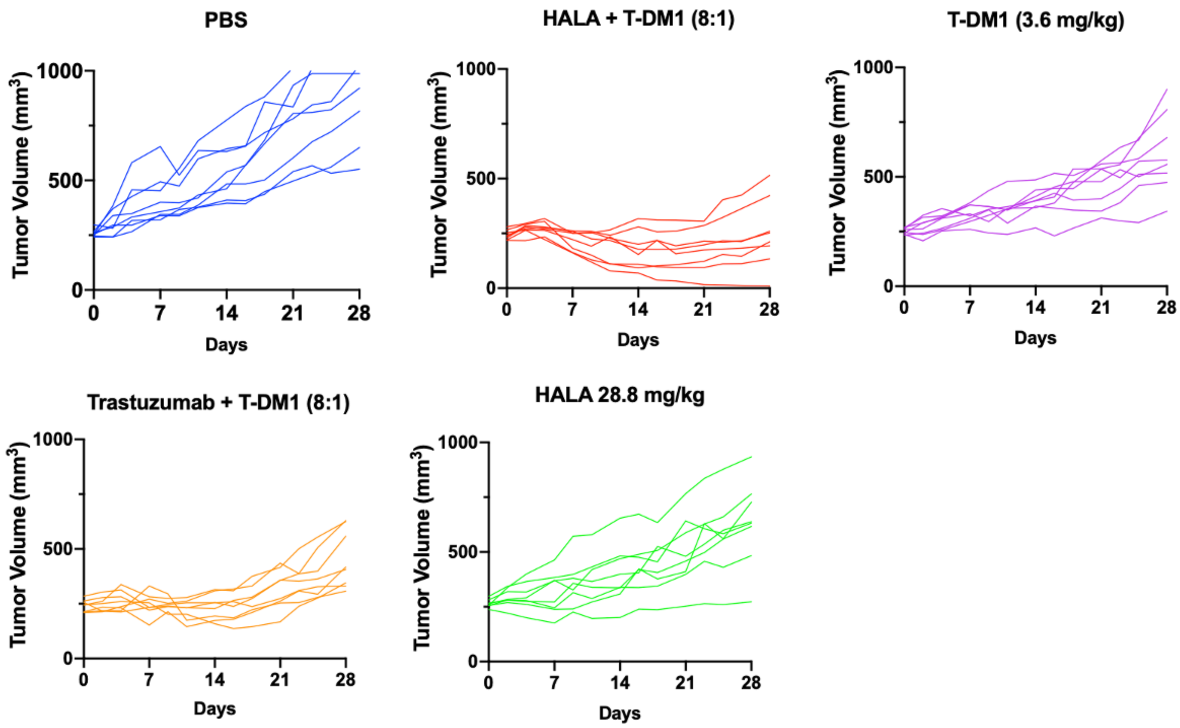


**Figure S6: NCI-N87 T-DM1 Efficacy**. Individual spider plots of T-DM1 efficacy in a high HER2 expression cell line (n=8 per group) with PBS control, 32.2 mg/kg HALA and 3.6 mg/kg T-DM1 (8:1), 3.6 mg/kg T-DM1, 32.2 mg/kg trastuzumab and 3.6 T-DM1, and 28.8 mg/kg HALA antibody only.

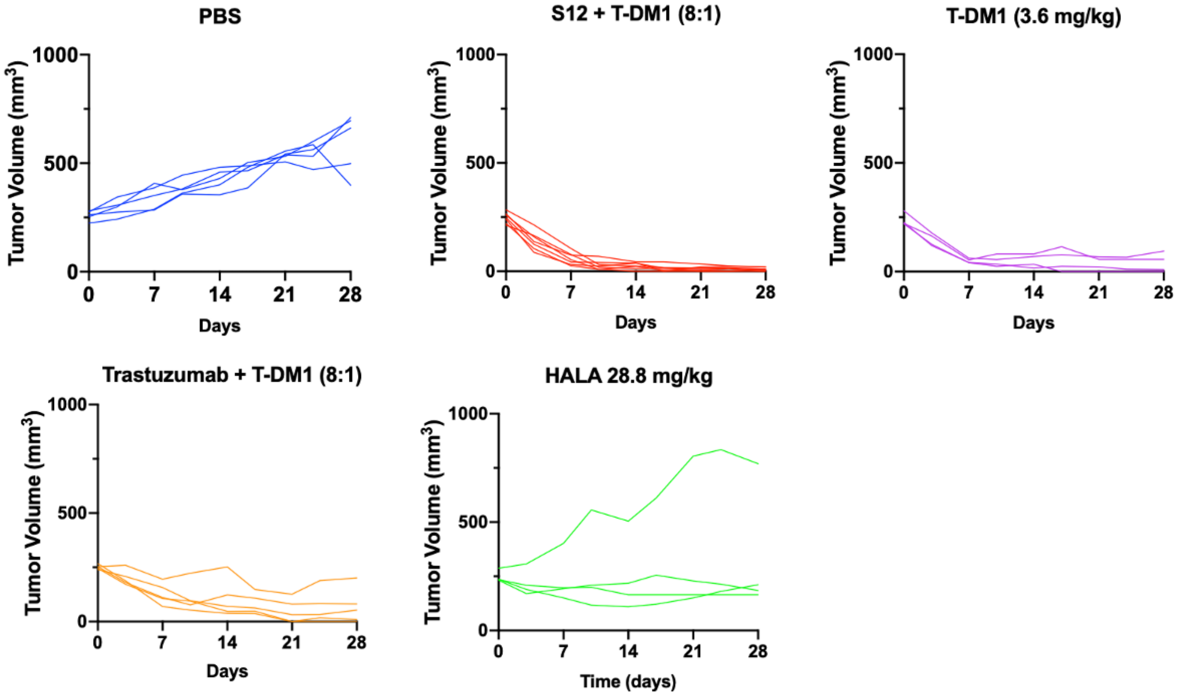


**Figure S7: MDA-MB-453 T-DM1 Efficacy**. Individual spider plots of T-DM1 efficacy in a low HER2 expression cell line with PBS control (n=5), 32.2 mg/kg HALA and 3.6 mg/kg T-DM1 (8:1) (n=5), 3.6 mg/kg T-DM1 (n=4), 32.2 mg/kg trastuzumab and 3.6 T-DM1 (n=5), and 28.8 mg/kg HALA antibody only (n=4).

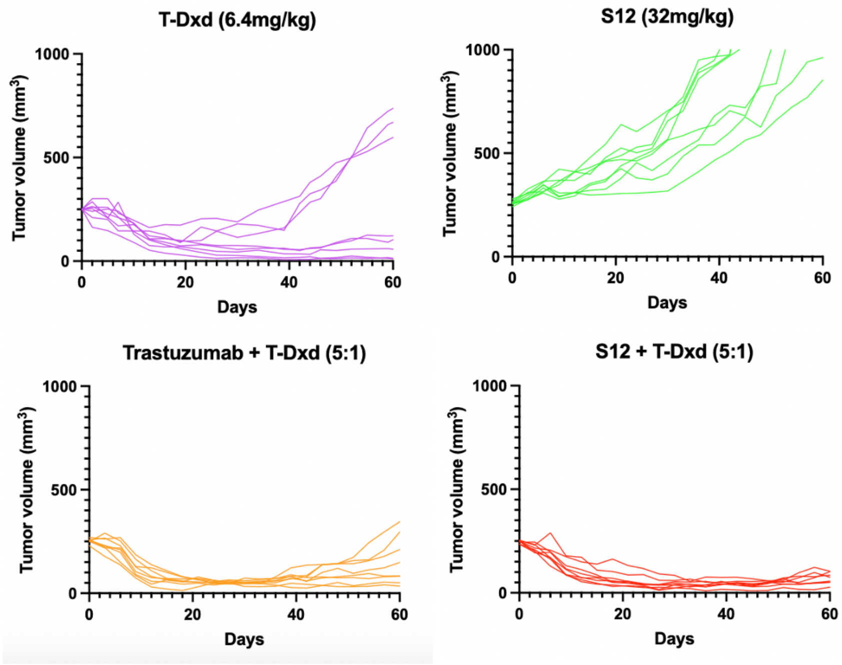


**Figure S8: NCI-N87 T-Dxd Efficacy**. Individual spider plots of T-Dxd efficacy in a high HER2 expression cell line with 6.4mg/kg T-Dxd, 32mg/kg HALA antibody only, 32mg/kg Trastuzumab and 6.4mg/kg T-Dxd, and 32mg/kg HALA antibody and 6.4mg/kg T-Dxd.

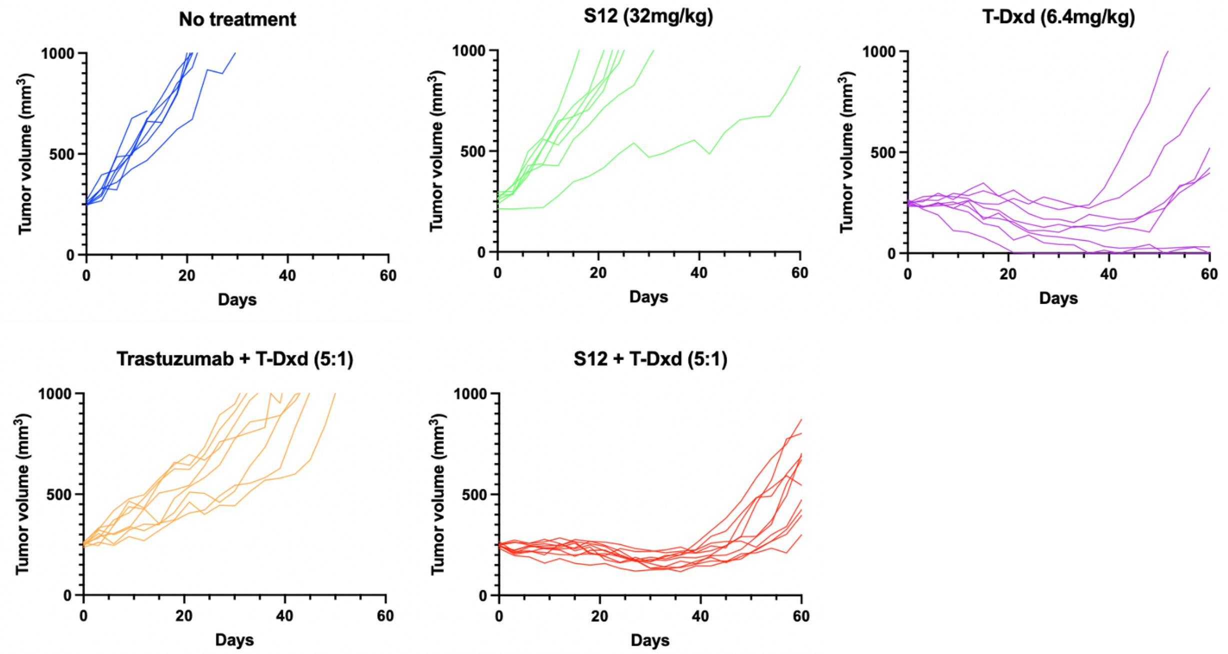


**Figure S9: CAPAN-1 T-Dxd Efficacy**. Individual spider plots of T-Dxd efficacy in a low HER2 expression cell line with no treatment control (n=7), 32 mg/kg HALA antibody only (n=7), 6.4 mg/kg T-Dxd (n=8), 32 mg/kg Trastuzumab and 6.4 mg/kg T-Dxd (5:1) (n=8), and 32 mg/kg HALA antibody and 6.4 T-Dxd (n=10).

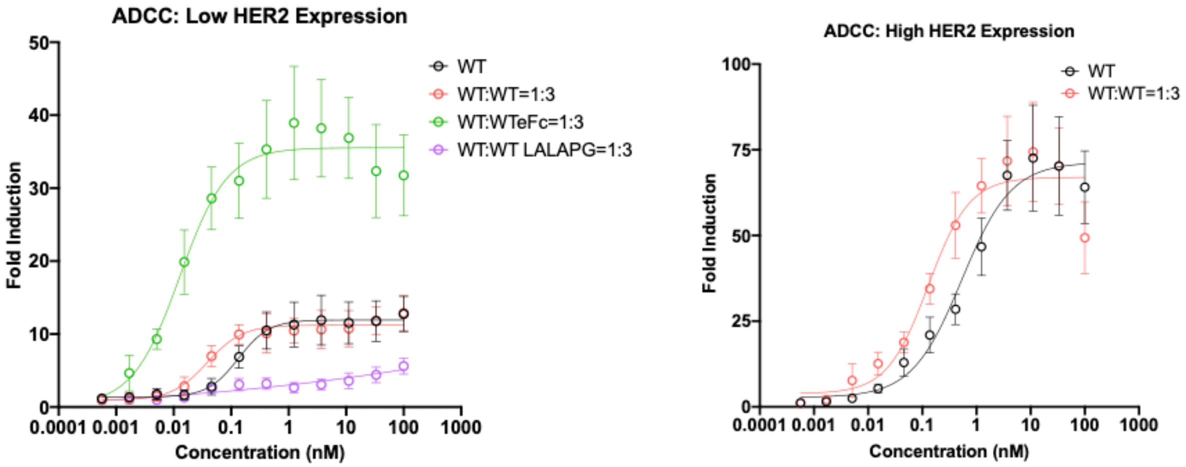


**Figure S10: Antibody Dependent Cellular Cytotoxicity (ADCC) of Antibody Combinations**. ADCC assays with WT carrier dose in the low expression cell line show increased ADCC with the Fc-enhanced trastuzumab (WTeFc) and low ADC activity when using an Fc-null carrier dose (WT LALAPG). The concentration on the x-axis is the WT antibody concentration independent of the carrier antibody. For example, a mixture of 1 nM WT antibody with 3 nM WT antibody carrier dose (WT:WT=1:3) is graphed as 1 nM. Therefore, a 1:3 WT:carrier dose slightly decreases the IC50 since the total antibody concentration is 4-fold higher. The WT and 1:3 WT:WT (low expression) and WT (high expression) data are duplicated from Figure 6 for comparison.

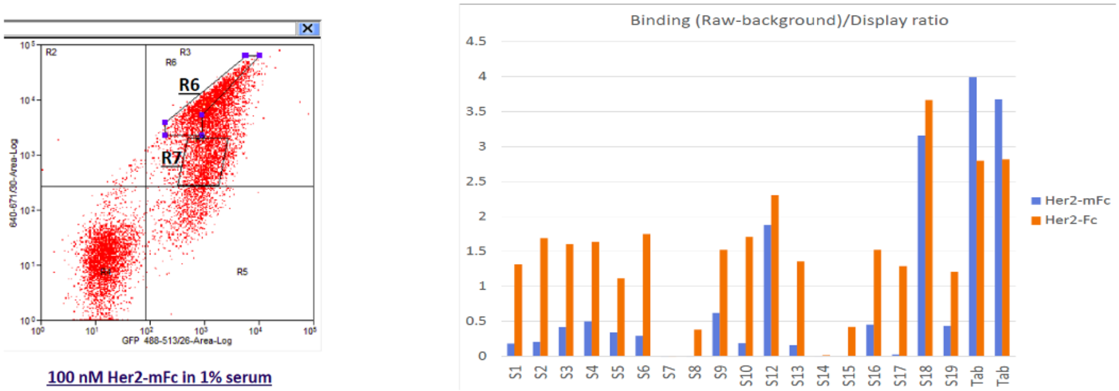


**Figure S11: Gating used for HALA sorting scheme and screening HALA candidates.** Variants were selected for further testing based on higher bivalent binding (HER2-Fc) compared to monovalent binding (HER2-mFc).

| Cell line | Measured HER2 Expression |
|---|---|
| MDA-MB-231 | $3.5 \times 10^4 \pm 320$ |
| Capan-1 | $2.36 \times 10^4 \pm 1{,}900$ |
| MDA-MB-453 | $3.76 \times 10^5 \pm 4{,}200$ |
| HCC-1954 | $4.4 \times 10^6 \pm 37{,}000$ |

Supplementary Table 1: Measured HER2 expression level of cell lines (Mean and SD).
